# Accelerated genome shuffling associated with rapid evolution of sexual conflict in seed beetles

**DOI:** 10.1101/2025.02.24.639998

**Authors:** Weiyao Chen, Jinshuai Zhao, Miaojin Lin, Changyou Liu, Liqiang Xie, Lixia Wang, Xiaoming Zhang, Yi Liao, Jinfeng Chen

**Affiliations:** State Key Laboratory of Integrated Management of Pest Insects and Rodents, Institute of Zoology, Chinese Academy of Sciences, Beijing 100101, China; University of Chinese Academy of Sciences, Beijing 100049, China; Institute of Cereal and Oil Crops, Hebei Academy of Agricultural and Forestry Sciences/Hebei Laboratory of Crop Genetics and Breeding, Shijiazhuang 050035, China; Institute of Crop Sciences, Chinese Academy of Agricultural Sciences, Beijing 100081, China; College of Horticulture, South China Agricultural University, Guangzhou, Guangdong 510642, China

**Keywords:** structural variation, Bruchinae, sex chromosome evolution, sex biased gene, sexual conflict

## Abstract

Structural variation (SV) has an extensive impact on the evolution of the genome. The diverse SV provide a material basis for genome evolution, which can affect gene expression, 3D genome structure, and even changes in complex chromosomal features through regional recombination, copy number variation, and chromosome rearrangement. Here, we assemble and compare the chromosome-level assemblies of three species (*Acanthoscelides obtectus*, *Callosobruchus chinensis*, and *C. maculatus*) in Bruchinae, a clade of Coleoptera that is often used as models to study sexual conflict, to investigate the influence of SV on the changes of various genomic characteristics during species evolution. Through comparative genomic analysis of Coleoptera species, we reveal extensive structural variants (SVs) within the Bruchinae family, which is linked to the ongoing activity of LINE transposons. We show that SV drive the rapid evolution of sex chromosomes and correlated with dynamic evolution of sex-biased gene expression in seed beetle species. Our analysis highlights the acquisition of novel genes on the Y chromosome through frequent gene translocations, gradually shaping them into genes exhibiting male-biased expression patterns. This underscores the Y chromosome’s role in promoting sexual dimorphism evolution. Our findings reveal the significant contributions of SV to genome evolution, specifically through their influence on sex chromosome evolution and the evolutionary dynamics of sex-biased genes.

## Introduction

Genomic evolution is a fundamental aspect of species evolution, documenting diverse information such as species diversity, variation, and adaptive changes throughout evolutionary processes (Ellegren and Sheldon 2008; Barrett and Hoekstra 2011; Barrett and Schluter 2008). It encompasses multiple facets, including changes in genome size, gene rearrangements, variations in gene sequences, regulatory changes in gene expression, as well as alterations in gene function and structure (Carvalho and Lupski 2016; Feuk, Carson, and Scherer 2006; Hastings et al. 2009; Lupiáñez et al. 2015; Pfeiffer, Goedecke, and Obe 2000). These molecular changes not only influence the genetic composition of populations but also play a crucial role in their ecological interactions and adaptation to dynamic environments (Jones et al. 2012). The rapid evolution of the genome underpins the emergence of numerous new species and traits (Davidson and Moczek 2024; Yoshida et al. 2023), thus constituting a pivotal focus in the study of species evolution.

Structural variation (SV) is a significant driving force in genome evolution, fostering evolutionary changes through diverse mechanisms. Firstly, SV can entail changes in chromosome numbers, such as deletions, duplications, insertions, or inversions, potentially impacting genome stability and function (Conrad et al. 2010; Huang et al. 2022). Secondly, they may alter gene sequence or position on chromosomes, thereby influencing gene expression patterns and regulation (Stranger et al. 2007; Gilbertson et al. 2022). Consequently, this can induce dysregulation, enhancement, or inhibition of gene function, shaping individuals’ phenotypic traits. Moreover, chromosomal SV may provoke imbalanced recombination between chromosomes, leading to chromosomal instability and aberrant genetic phenomena like chromosome breakage and rearrangement (Aguilera and Gómez-González 2008; Li and Heyer 2008). In terms of gene regulation, SV can disrupt the original Topologically Associated Domain (TAD) structure of the genome, influencing the spatial proximity between genes and their regulatory elements, thereby indirectly altering gene expression (Gilbertson et al. 2022; Rocks et al. 2022). Sex chromosome sequence disparities constitute a prominent feature underlying sex differences, and variations in sex chromosomes can contribute to alterations in sexual dimorphism (Kaufmann et al. 2023; Graves 2016). SV influences the evolution of sex chromosomes through mechanisms such as chromosome recombination and the translocation of sex-linked genes, thereby contributing to the development of sex-specific traits (Wang et al. 2022; Tosto et al. 2023). Furthermore, SV plays a pivotal role in facilitating the emergence of novel traits. Variations such as single nucleotide polymorphisms (SNPs), sequence insertions, and deletions can impact the genotypes of key regulatory genes across individuals, thereby influencing the expression of corresponding traits (Montgomery et al. 2013). Minor mutations in protein sequences can alter their overall structure, introducing new functions and interaction patterns (Li et al. 2024; Anzelon et al. 2021). Additionally, new genes may arise from sequence insertions, duplications, or transposon variations, contributing novel functions or expression patterns (Long et al. 2003; Chen, Krinsky, and Long 2013).

Under natural selection, sexual dimorphism arises from sexual conflict at loci, which can be mitigated through gene translocation and sex-biased expression (Sayadi et al. 2019; Tosto et al. 2023). In many species that engage in sexual reproduction, females and males invest different resources in many biological processes, including metabolism, reproduction, and offspring rearing, and this antagonism serves as the precursor for the emergence of sexual conflict (Harrison et al. 2015; Han et al. 2022). Sex chromosomes serve as prime loci for mitigating sexual conflict through gene translocation, facilitated by inherent sex-specific differences (Kaufmann et al. 2023; Catalán, Macias-Muñoz, and Briscoe 2018). Many studies have found that in XY systems, the X chromosome is enriched with more female-biased genes (Lasne et al. 2023), while in ZW systems, the Z chromosome is enriched with more male-biased genes (Catalán, Macias-Muñoz, and Briscoe 2018). The uniqueness of the Y chromosome in male individuals also provides a possibility for the genes located on it to assist in mitigating sexual conflict. A recent study has found that the transfer of a gene to the Y chromosome can help alleviate conflicts related to sexual dimorphism in body size, indicating the significant contribution of the Y chromosome in resolving sexual conflict (Kaufmann et al. 2023). However, the lack of recombination and enrichment of sequence mutations on the Y chromosome (Charlesworth and Charlesworth 2000) often results in its scarcity of genes and abundance of transposons, making the assembly of the Y chromosome extremely difficult, thus impeding research into the relationship between Y chromosome genes and sex-biased genes.

Seed beetles are a classic model for studying sexual selection and sexual conflict, and they have a typical XY sex-determination system (Sayadi et al. 2019). There are notable trait variations among different seed beetles, encompassing diverse dietary habits, physical attributes, and sex pheromone expression (Wilson et al. 1997; Savalli and Fox 1999; Salehialavi, Fritzsche, and Arnqvist 2011; Fritzsche, Booksmythe, and Arnqvist 2016). Moreover, distinct sexual dimorphisms are observed across various seed beetle species, implying potential differences in sexual conflict. For instance, in *Callosobruchus chinensis*, antennal morphology differs between males and females, whereas in *C. maculatus*, no such distinction in antennal shape exists between the sexes (Hu, Zhang, and Wang 2009). The seed beetles *C. maculatus* are considered as a classic model for studying sexual conflict, including reproductive characteristics (Hotzy et al. 2012) and body size differences (Berger et al. 2016). However, due to the current scarcity of comprehensive genome resources in seed beetles (Kaufmann et al. 2023; Immonen et al. 2023; Lu et al. 2024; Arnqvist et al. 2024), studying the role of SV in seed beetles’ traits and sexual conflict through comparative genomics methods remains challenging.

In this study, we assembled the chromosome-level genomes of three seed beetle species: *Acanthoscelides obtectus*, *C. chinensis*, and *C. maculatus*. Utilizing whole-genome sequencing data from both male and female samples, we successfully identified their X and Y chromosome. Comparative analyses revealed a striking landscape of extensive and large-scale genome rearrangements within the Bruchinae family. Through comparative analysis of Bruchinae family genomes, this study reveals that SV significantly influenced genome evolution across multiple dimensions, including sex chromosome differentiation and sex-related genetic conflict. These findings underscore the role of SV in driving species evolution, enhance our understanding of genetic mechanisms shaping biological diversity.

## Result

### *De novo* genome assembly of three diverse seed beetle species

To advance our understanding of genome evolution in seed beetle species and enable comparative genomic analyses of previously inaccessible regions (e.g., the Y chromosome), we generated *de novo* genome assemblies for three diverse seed beetle species using long-read sequencing data. The first species we selected for sequencing is *C. maculatus*, which is served as a model species for studying insect evolution and ecology and has previously been assembled at the contig level (Sayadi et al. 2019). We generated 2.5 million High fidelity (HiFi) reads, resulting in a total of 44.6 Gb of raw sequencing data, corresponding to approximately 38-fold coverage of its estimated genome size 1,107 Mb. We assembled the HiFi reads using hifiasm and obtained a consensus genome assembly of 1,153 Mb with contig N50 up to 13.1 Mb. Additionally, we *de novo* assembled the genomes of two other seed beetle species, *C. chinensis* and *A. obtectus*, to expand our understanding of the genome evolution within the Bruchinae family. For these two species, we generated 21.3 and 1.3 million Oxford nanopore Technology (ONT) ultra-long reads (90.6 Gb and 25.7 Gb), corresponding to approximately 143-fold and 28-fold genome coverage of their estimated genome sizes (642 Mb and 866 Mb), respectively. We applied NextDenovo to assemble the ONT long reads, resulting in assemblies of 633 Mb for *C. chinensis* and 917 Mb for *A. obtectus*, with contig N50 of 3.6 Mb and 4.2 Mb, respectively. These contig-level assemblies were further polished with paired-end short reads and anchored using Hi-C interaction reads to generate the chromosome-scale genome assemblies of the three species. Approximately 99.5%, 91.6%, and 99.36% of the assembled contigs are placed into eleven pseudo-chromosomes for seed beetles *C. chinensis*, *C. maculatus*, and *A. obtectus*, respectively (Figure 1A-B and S1A).

**Figure 1.**
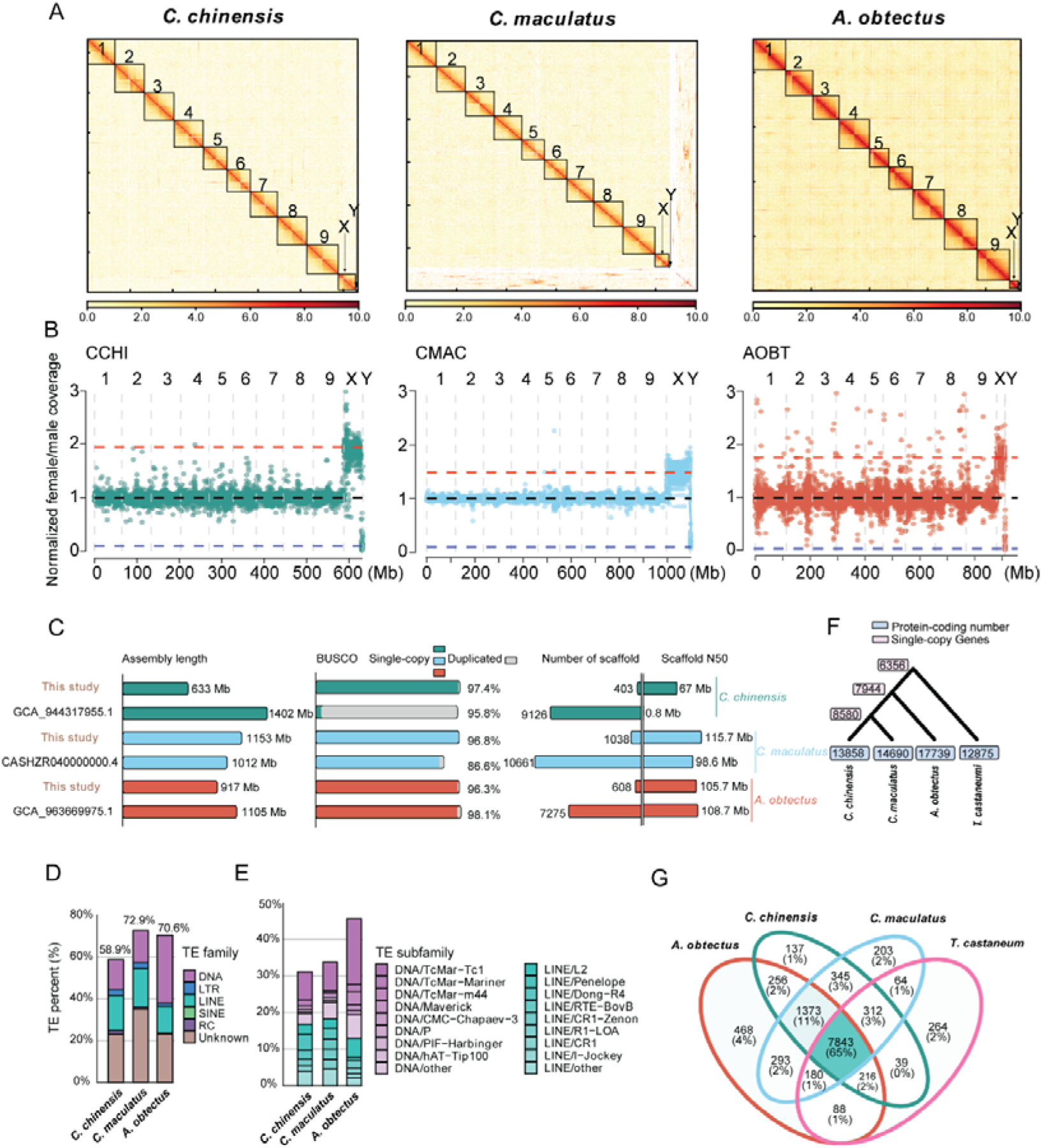
Genome assembly and annotation of *C. chinensis*, *C. maculatus*, and *A. obtectus*. **A,** Hi-C contact maps of the *C. chinensis*, *C. maculatus*, and *A. obtectus* genome assembly. Black boxes denote putative chromosomes. The white indicates the weakest scaffold interaction, while the red indicates the strongest interaction. **B,** Normalized female/male whole genome sequencing coverage over the three seed beetle species genome assembly. Each point represents a 100kb window. Blue, black, and red horizontal lines indicate expected coverage over Y-linked, autosomal, and X-linked scaffolds, respectively. **C,** Comparison of genome assembly parameters with previously published genomes for the three seed beetle species. **D,** Repeat landscape of the three seed beetle species genomes. **E,** Subfamily of DNA transposons and LINE transposons landscape of the three seed beetle species genomes. **F,** Phylogenetic trees and gene number of three seed beetle species and the model species *T. castaneum*. **G,** Comparative analysis of orthologous families from the three seed beetle species and the model species *T. castaneum*.

During the preparation of this manuscript, genome assemblies of the three seed beetles are published or released (Arnqvist et al. 2024; Immonen et al. 2023; Lu et al. 2024; Papachristos, Sayadi, and Arnqvist 2024). Compared to the previous version of *C. chinensis* (Papachristos, Sayadi, and Arnqvist 2024), the number of contigs in current assembly has been drastically reduced from 9,126 to 403, and the contig N50 has significantly increased from to 0.8 Mb to 67 Mb. Furthermore, the genome size has been reduced from 701 Mb to 633 Mb by removing the redundancy between haplotypes in the current assembly. The current assembly of *C. maculatus* is comparable to the newest genome of *C. maculatus* (Lu et al. 2024) (Figure 1C and Table S1). Similarly, the current assembly of *A. obtectus* is comparable to the previous version (Immonen et al. 2023) in term of genome size and contig N50, however demonstrates improved contiguity (Figure 1C). The new genome assemblies of *C. chinensis*, *C. maculatus*, and *A. obtectus* also demonstrate high completeness in gene space, containing 97.4%, 96.8% and 96.3% of conserved Insecta benchmarking universal single-copy orthologs (BUSCO) genes (n=1,367) as complete single copies, respectively (Figure 1C and Table S1).

### Identification of seed beetles’ sex chromosomes

To identify the sex chromosomes, we aligned the whole-genome sequencing data obtained from both female and male samples to the assembled pseudo-chromosomes and calculated the female-to-male coverage ratio of each bin with a 100kb sliding window. Cytological studies have indicated a correlation between the sex of seed beetles and the number of X and Y chromosomes (Angus et al. 2011), where females possess two X chromosomes and males have one X chromosome paired with one Y chromosome, resulting in an expected coverage ratio of X chromosome to autosomes of approximately 2:1 in females. In contrast, the ratio involving Y chromosome should approach zero, as it is present only in males. Consistent with cytological analysis, we observed a specific chromosome in each seed beetle exhibits the expected coverage characteristic of an X chromosome along its entire length (Figure 1B). Comparing the identified X chromosome of the *C. maculatus* with the previously published X chromosome (Lu et al. 2024; Arnqvist et al. 2024), we discovered that they share homologous relationships (Figure S2). Furthermore, these X chromosomes from the three seed beetles possess unique Hi-C matrices and exhibit sequence homology with each other (Figure S3), confirming that they are X chromosomes.

We found that the shortest pseudo-chromosome in each seed beetle has the expected coverage pattern of Y chromosome (Figure 1B). Each of them is composed of several contigs, and the presence of Hi-C signals between them results in the formation of an independent Hi-C matrix unit, signifying a segment within the Y chromosome (Figure S4 A-C). Building on previous studies that identified several Y-link contigs in *C. maculatus*, we conducted a comparative analysis to assess the homology between the published Y-linked contigs and the Y chromosome sequence we assembled in *C. maculatus.* A high level of sequence conservation was observed between the identified Y chromosome and the previously published Y-linked contigs of *C. maculatus*, confirming the accuracy of our Y chromosome identification (Figure S4 D-F). However, the Y chromosome of the current genomes appears to be relatively smaller in size compared to the Y-linked contigs reported in other studies (Kaufmann et al. 2023). Based on previous comparisons of sex chromosomes across various populations of seed beetles (Angus et al. 2011), we hypothesized that this discrepancy could be attributed to population-specific genetic variations.

In summary, we identified the X and Y chromosomes in three seed beetles. For the X chromosome of *C. maculatus*, we assembled an X chromosome of similar size to the previous version (Lu et al. 2024) through 11 contigs (56.0 Mb vs. 47.3 Mb). As for *A. obtectus* and *C. chinensis*, we produced the first chromosome-level X chromosomes (29.8 Mb and 40.7 Mb). Regarding the Y chromosome, the previous version of *C. maculatus* (Sayadi et al. 2019; Kaufmann et al. 2023) generated Y-linked contig fragments, whereas we utilized Hi-C signals to assemble a scaffold-level Y chromosome (3.9 Mb). Similarly, for *A. obtectus* and *C. chinensis*, we also assembled scaffold-level Y chromosomes (1.9 Mb and 3.6 Mb) (Table S1).

### Genome annotation of three diverse seed beetle species

Compared to genomes of other Coleoptera species, a notable distinctive feature of the seed beetle genomes is their significantly higher content of repetitive sequences. Specifically, 58.9%, 72.9%, and 70.7% of the genomes in *C. chinensis*, *C. maculatus*, and *A. obtectus,* respectively, consist of repetitive sequences (Figure 1D and Tables S2-S4). In *A. obtectus*, DNA-type transposons (32.72%) constitute the largest proportion among all transposable element (TE) types in the genome (Table S4). In contrast, both DNA-type (14.28% and 16.04%) and LINE (16.85% and 18.93%) TEs are prominent in *C. chinensis* and *C. maculatus* (Figure 1E, Tables S2-S3). This suggests that a lineage-specific amplification of LINEs may have occurred in the ancestral lineage leading to both *C. chinensis* and *C. maculatus*. The most abundant superfamily among DNA-type transposons is Tc1/*Mariner*, which accounts for 7.88% to 20.08 % of the genomes of the three seed beetle species. The abundance of the LINE retrotransposons shows a relatively consistent distribution across superfamilies and species (Figure 1E).

The predicted number of protein-coding genes is 13,858 for *C. chinensis,* 14,690 for *C. maculatus*, and 17,739 for *A. obtectus*, with average CDS length ranging between 1,297 and 1,529 bp (Figure 1F and S1B-C). Protein-coding genes that have functional annotations make up 72.7%, 71.5%, and 68.9% of the total predicted genes, respectively. The completeness of the gene sets for the three seed beetle species was assessed using the insecta BUSCO gene set (n=1,367), with 93.8% to 95.2% of these genes identified as complete (Table S1). Comparing the predicted genes with *Tribolium castaneum* genome (Kim et al. 2010) (Figure 1G), we found a total of 7,843 (65%) orthologous families shared across all four species, and 1,373 (11%) orthologous families specific to the Bruchinea family. Additionally, we identified 7,944 strict single-copy genes among three seed beetle species, of them, 6,356 are also shared with the *T. castaneum* (Figure 1G). In summary, we generated high-quality gene annotations for these three seed beetle species and demonstrated that over half of the genes are conserved between Bruchinae and *Tribolium castaneum*.

### Extensive chromosomal rearrangements during the evolution of species in Bruchinae family

To explore the genome evolution of these three seed beetle species, we extended the comparative genomic analysis to eight species with varying divergence time within the Coleoptera family: *Pachyrhynchus sulphureomaculatus* (Van Dam et al. 2021), *Pyrochroa serraticornis* (GCA_963669975.1), *T. castaneum* (Kim et al. 2010), *T. freemani* (GCA_939628115.1), *T. confusum* (GCA_029207805.1), *Propylea japonica* (Zhang et al. 2020), *Rhagonycha fulva* (GCA_905340355.1), and *Pogonus chalceus* (Ando et al. 2018), using *Drosophila melanogaster* as the outgroup. A maximum likelihood phylogenic analysis based on 1,397 single-copy orthologous genes allowed us to estimate that *C. chinensis* and *C. maculatus* split from their common ancestor approximately 30 million years ago (Mya), while *A. obtectus* diverged from the *Callosobruchus* genus around 56 Mya (Figure 2A). The genome-wide synteny analysis across these species confirms the homologous relationships of chromosomes and karyotypes among most Coleoptera insects (Figure 2B), which is consistent with observations from earlier studies (Van Dam et al. 2021). It also establishes that the genomes of the three seed beetle species in the Bruchinae family retain well-conserved synteny in both autosome and X chromosome (Figure 2B). However, extensive large-scale chromosomal rearrangements were observed between the Bruchinae species and other Coleoptera species, except for the X chromosome, which exhibited conserved gene collinearity in all Coleoptera species examined (Figure 2B and S5A-E). For example, *P. sulphureomaculatus*, the phylogenetically closest species to Bruchinae in our study, displays conserved karyotype with most Coleoptera species, except for those in Bruchinae (Figure S5A-E, H). We found that most genes located on the autosome of *A. obtectus* have homologs in the genomes of all other Coleoptera species, but their genomic positions have changed (Figure S5F-G). These findings reveal that gene synteny has been largely decayed between Bruchinae species and other Coleoptera species, despite the fact that most genes remain present in both.

**Figure 2.**
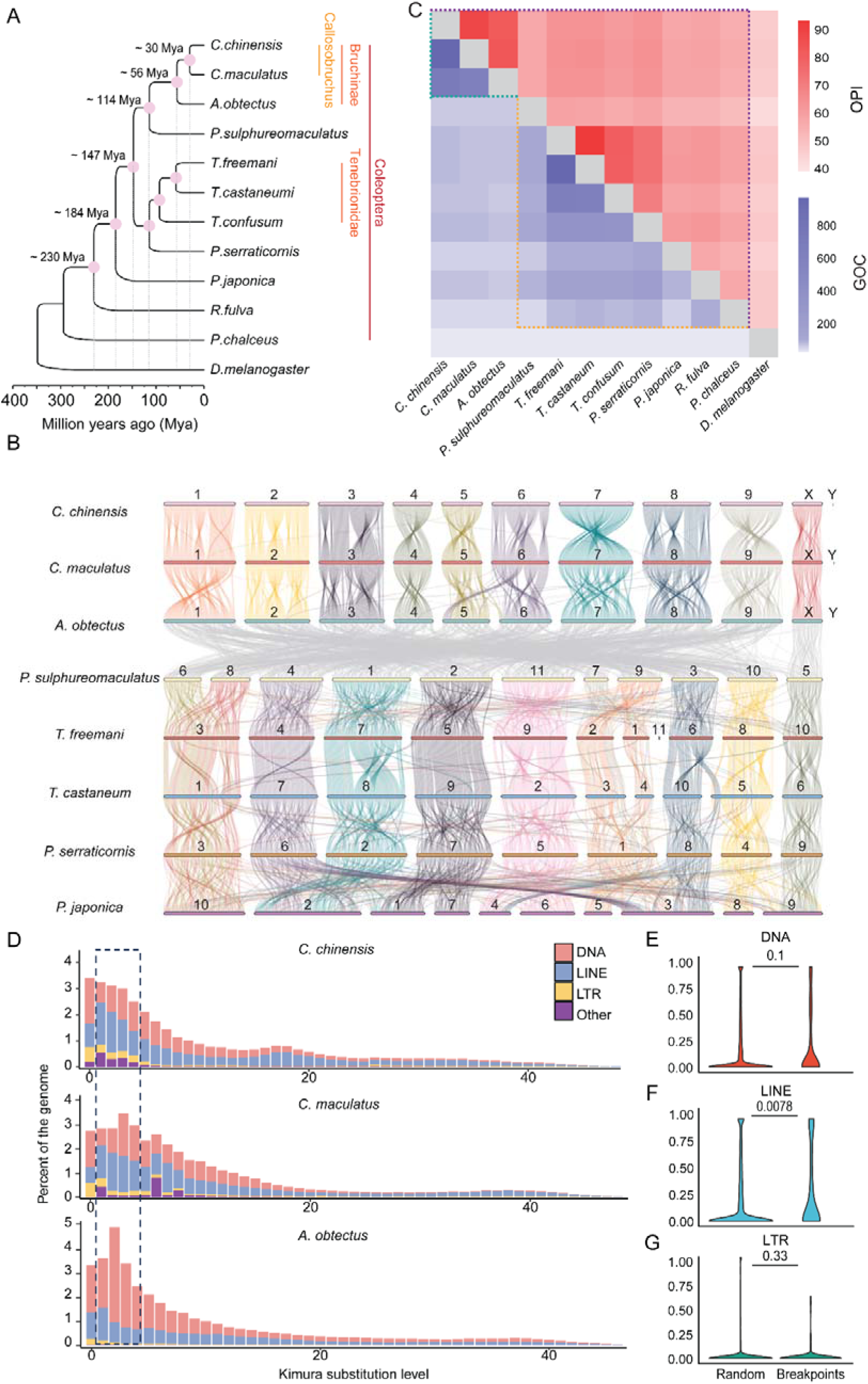
Widespread genome recombination in Bruchinae. **A,** The phylogenetic tree of different species in the Coleoptera. A maximum likelihood tree of different Coleoptera species with *D. melanogaster* as outgroup was established by iqtree. **B,** Pairwise whole-genome alignments across 8 Coleoptera genomes. Chromosome IDs of each species are labeled on chromosomes. The color of the collinear region among *P. sulphureomaculatus*, *T. freeman*, *T. castaneum*, *P. serraticornis*, and *P. japonica* are referenced by *P. sulphureomaculatus*. The color of the collinear region among *C. chinensis, C. maculatus* and *A. obtectus* are referenced by *C. chinensis*. **C,** A heat map matrix for orthologous proteins identity (OPI) and gene order conservation score (GOC) among Coleoptera and Bruchinae. The difference boundary between identity and GOC along evolutionary distance is highlighted with dashed lines. **D,** Insertion history of TEs among seed beetles. Vertical bars show the frequency of TE insertions during the evolution of species. Recent DNA and LINE transposon amplification are highlighted by dashed lines. **E-G,** Contents of DNA, LINE and LTR in rearranged breakpoint regions and random regions. Breakpoint are identified between our assembly and Lu *et al*. genome. The breakpoint with 1k flanks region are compared with random regions of the same number and length of genomes. The number on the horizontal line above each two boxes represents *p*-values (two-sided *t*-test).

To quantitatively describe the extent of chromosomal rearrangements in Bruchinae species, we computed orthologous proteins identity (OPI) (Table S5) and the Gene Order Conservations (GOC) (Van Dam et al. 2021) scores (Table S6) across all pairwise comparisons among the eleven Coleoptera species using highly conserved gene set from BUSCO (version 4; insecta BUSCO gene, n = 1367) (Figure 2C). The results show that *A. obtectus* has the highest OPI with *C. maculatus* (84.2%) and *C. chinensis* (83.1%) compared to other Coleoptera species, which have OPI values ranging from 46% to 65%. Similarly, the Bruchinae species have the higher level of GOC scores (624-953) compared to other Coleoptera species (10-65) (Figure 2C). Both results demonstrate that the Bruchinae species have experienced an extensive decay of synteny since their divergence from other Coleoptera species. We also compared sequence similarity of syntenic segments between Tenebrionidae and Bruchinae species, taking their differentiation times into consideration, and observed that species in Bruchinae exhibited comparatively higher sequence divergence. For example, the sequence divergence between *A. obtectus* and *C. chinensis* (∼56MYA, 87.45%) is notably higher than that observed between *T. castaneum* and *T. freemani* (∼58MYA, 90.42%). Even between *C. chinensis* and *C. maculatus* (∼30MYA, 89.75%), the sequence differences are greater than those found within the Tenebrionidae, indicating a more rapid sequence divergence within the Bruchinae family (Figure S6A-C).

The activity of TEs is a major mechanism for creating structural variants (SVs) (Cordaux and Batzer 2009; Beck et al. 2010). We then investigated whether any TE families are associated with the rapid occurrence of chromosomal rearrangements in Bruchinae species. We observed a lineage-specific recent burst of LINE retrotransposons in the three seed beetle species, while the genomes of *A. obtectus* and most other Coleoptera species have experienced greater amplification of DNA transposons compared to LINE retrotransposons (Figure 2D and Figure S7). By comparing two available *C. maculatus* genomes (Figure S2D), we found that the LINE retrotransposons were significantly enriched at regions where SVs breakpoint occur (Figure 2E-G). These results suggest that the lineage-specific amplification of LINEs may contribute to the higher rate of chromosomal rearrangements in Bruchinae species compared to other Coleoptera species.

### The impact of SV on the evolution of sex chromosomes in the Bruchinae family

SV drive the evolution of sex chromosomes through various mechanisms, including genome recombination, gene traffic, and chromosome fusion (Wang et al. 2022; Gu et al. 2019). To explore the sex chromosome evolution in the Bruchinae family, we performed comparative sequence analysis of the X and Y chromosome between species of different divergence within this family. Synteny analysis shows that the synteny of the X chromosome can be largely maintained across most of families within the Coleoptera order (Figure 2B). For example, approximately 59.4%-65.9% of genes located on the X chromosome of *T. castaneum* have orthologous counterparts on the X chromosomes of three seed beetle species, whereas the proportion on autosomes is only 8.3%-9.9% on average (Figure 3A). Similar results are also observed when comparing *P. sulphureomaculatus* to species in Bruchinae (Figure S8). The comparative genomic analysis between species within the Bruchinae family also observed extensive SVs on the X chromosome (Figure 3B) but most of the X-linked genes have their orthologs on the X chromosome of at least one of the other species (65.4%, 66.9% and 72.0% for *A. obtectus*, *C. chinensis* and *C. maculatus,* respectively) (Figure 3C). These results reveal that, despite extensive SVs occurred on the X chromosome of species in the Bruchinae family, gene content and synteny retain largely conserved, likely due to the evolutionary functional constraints of X-linked genes and their chromosomal locations (Mathers et al. 2021).

**Figure 3.**
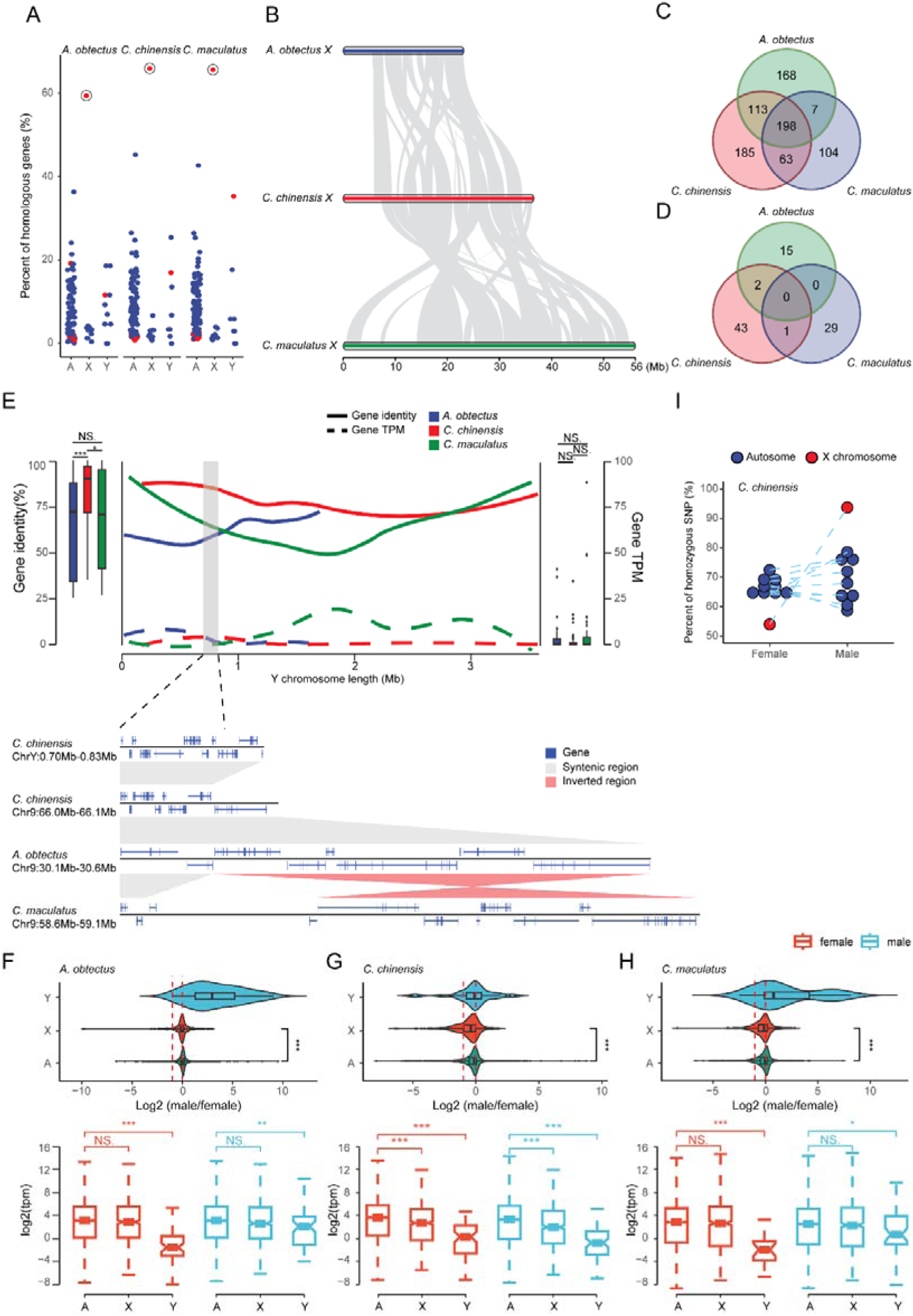
Evolution of Sex chromosomes in Bruchinae. **A,** Chromosomal conservation of orthologous genes between Bruchinae and *T. castaneum*. Each point represents the percentage of orthologous genes between different chromosomes of the *T. castaneum* and the autosomes, X chromosome, and Y chromosome of the corresponding Bruchinae species, relative to the total number of genes on each chromosome of the *T. castaneum*. The X chromosome of *T. castaneum* is marked with red dots, which exhibit the highest percentage of orthologous genes with Bruchinae species’ X chromosome. **B,** Widespread chromosomal recombination between the X chromosome of the seed beetles. Gray blocks represent regions of gene collinearity. **C-D,** Conservatism of orthologs between seed beetles’ X (**C**) and Y (**D**)chromosome. **E,** Sequence characteristics and gene expression of Y chromosome. The overall similarity between Y chromosome genes and their orthologs in autosomes is depicted by box plot. The distribution of gene identity (solid line) and expression levels (dashed line) along the Y chromosome is illustrated by a line graph. The collinearity of a segment of *C. chinensis*’ Y chromosome with autosomes is magnified at the bottom. **F-H,** Dose expression pattern of sex chromosomes for *A. obtectus* (**F**), *C. chinensis* (**G**) and *C. maculatus* (**H**) in head. The log2 of the male-to-female expression ratio for autosomal and sex-linked genes is depicted in the violin plot. Expression of autosomal and sex-linked genes in males and females is depicted in the box plot. Female samples are represented by red and male samples are represented by blue. *P* values (\**P* < 0.05, \*\**P* < 0.01, \*\*\**P* < 0.001) were derived from two-side Wilcoxon matched-pairs signed-rank tests. **I,** Allelic heterozygosity in *C. chinensis*. Each point represented a chromosome and X chromosome is represented by red dot.

Conversely, the Y chromosome exhibit very little conservation in both gene content and synteny. Among species in the Bruchinae family, the Y chromosome shows minimal synteny conservation and lack gene content conservation altogether (Figure 3D). This dramatic difference on sequence content may be attributed to the high rate of sequence gain and loss during the evolution of Y chromosome as a result of specific SV events. For example, we identified a genomic segment spanning ∼130kb and containing 14 genes on the Y chromosome of *C. chinensis* that was derived from chromosome 9, which remain synteny across all three species in Bruchinae (Figure 3E). These results suggest that gene-related SVs lead to the rapid turnover of gene content on the Y chromosome of species in Bruchinae, consistent with observations in human and *Drosophila* (Vicoso 2019).

We next investigated to what extent SV can affect the expression of sex-linked genes by comparing gene expression profiles between male and female tissues across the three seed beetle species (Figure 3F-H). At a genome-wide scale, our analysis of orthologs indicates no significant differences in gene expression levels on the X chromosome between male and female individuals in all three seed beetle species (Figure 3F-H). The ratio of X chromosome expression levels between males and females was comparable to that of autosomes (Figure 3F-H), indicating that the gene expression on the X chromosome in three species is compensated in males to reach levels similar to autosomes. This is demonstrated by further analyzing RNA-Seq data from male and female individuals where we found that males exhibit a higher proportion of homozygous X chromosome transcripts, while females show heterozygous X chromosome transcripts, comparable to the autosomes, in *C. chinensis* (Figure 3I and Figure S9). This result suggests that X chromosome dosage compensation is achieved by upregulating the expression level of the X-linked genes in males. In contrast, the expression level of orthologs on the Y chromosome is significantly lower than those on autosomes, even in the head tissues of male individuals (Figure 3F-H). This discrepancy between the X and Y chromosomes may be due to the rapid evolutionary turnover on the Y chromosome, leading to incomplete and unstable gene expression levels. A similar pattern was also observed in the gut and gonad tissues (Figure S10). Furthermore, we found that genes with shorter evolutionary histories tend to exhibit lower expression levels (Figure S11), where we observed the lowest gene expression levels and the most recent chromosomal rearrangements.

### SV drives the rapid evolution of sex-biased genes

The seed beetles, particularly *C. maculatus*, have been widely studied in evolutionary biology due to their pronounced sexual dimorphism and the sexual conflict that arises from their mating behavior (Kaufmann et al. 2023; Kaufmann et al. 2021; Sayadi et al. 2019; Hotzy et al. 2012). To investigate the genomic architecture underlying sexual conflict in seed beetles, we collected RNA-seq data from three tissues, mature gonads, guts, and heads, of adult males and females in *A. obtectus*, *C. chinensis*, and C. *maculatus*, and compared them to *T. castaneum*. In total, we identified 8367, 10005, 8882, and 8609 sex-biased genes (SBGs) across tissues in *T. castaneum, A. obtectus*, *C. chinensis*, and C. *maculatus*, respectively, with the majority (86.6-97.6%) predominantly observed in gonads (Figure 4A). We also identified a higher proportion of female-biased genes in head tissues from *C. chinensis* (13.0%) and *C. maculatus* (18.5%) compared to *A. obtetus* (2.1%), suggesting that lineage-specific transcriptome evolution related to sexual conflict may have emerged in the head tissue following their divergence from *A. obtectus*. Additionally, we found that the set of genes displaying sex-biased expression differ substantially between *C. chinensis* and *C. maculatus* (Figure 4B and Figure S12), with only 56.8% of the single copy genes being conserved between these two species in gonads. Furthermore, we observed that a substantial fraction of SBGs exhibits different directions of bias between species, even within the same tissue (13.9% in guts; 22.9% in heads; 64.3% in gonads). In other words, a gene may be male-biased in one species and female-biased in another. These results indicate that the expression pattern of sex-biased genes can evolve or turnover rapidly between seed beetle species.

**Figure 4.**
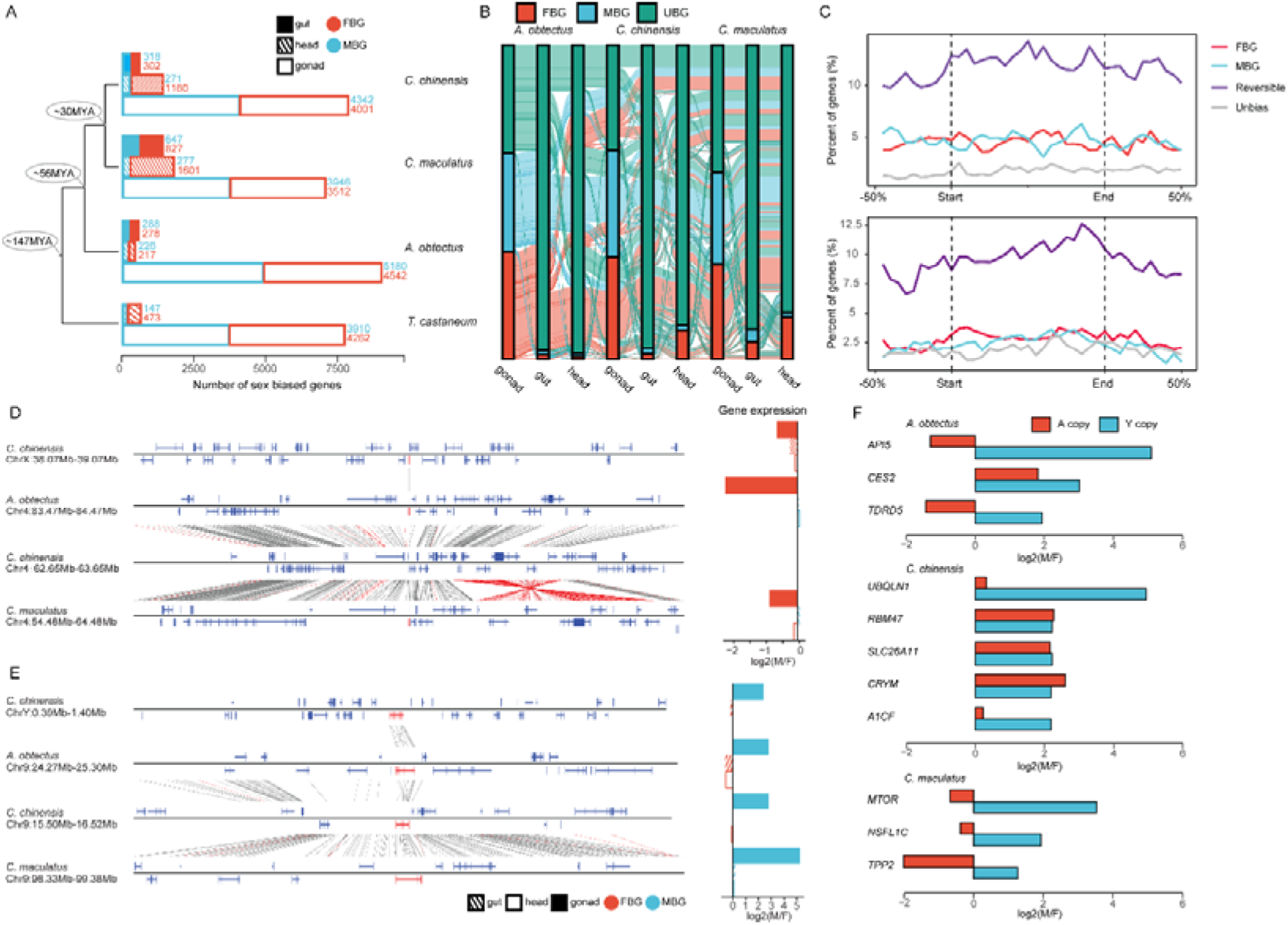
The evolution of sex-biased genes and the role of chromosomal recombination therein. **A,** Comparison of the number of sex biased gene between different species and tissues. Identify male bias genes (MBG) and female bias genes (FBG) through male and female samples from different tissues. The specific number of genes is indicated by numbers. **B,** The Sankey diagram showing dynamic variation of SBGs across different tissues and species. UBG, unbiased gene. **C,** Distribution of sex-biased gene density between *A. obtectus* and *C. chinensis* (top) or between *A. obtectus* and *C. maculatus* (bottom) along structural variation. ‘Reversible’ represents the orthologous with different sex biased expression patterns in different species. **D,** Non-replicating translocation event of gene. **E,** Replication translocation event of gene. The genes that undergo translocation are marked in red on the collinearity chart, and the corresponding sex-biased expression patterns in different tissues are shown in the bar plot. **F,** The expression of genes with strong sex bias on the Y chromosome and their orthologous genes on the autosomes. Expression of Y-linked genes are calculated by testis and ovary tissues.

We next sought to investigate if any genomic features that may be associated with changes of sex-biased genes in *C. chinensis* and *C. maculatus* relative to *A. obtectus*. We focused on large genomic variants and identified 270 (225 translocation and 45 inversion) and 237 (191 translocation and 46 inversion) SVs larger than 50 kb in *C. chinensis* and *C. maculatus*, respectively, relative to *A. obtectus*. We observed that sex-biased genes either identified between *C. chinensis* and *A. obtectus*, or between *C. maculatus* and *A. obtectus*, are significantly enriched in genomic regions that overlap with SVs (Figure 4C and Figure S13). This suggests that SVs may be a potential driver underlying rapid evolution of sex-biased genes in seed beetle species.

### Translocation-based gene birth on the Y chromosome of seed beetle species

Gene traffic has been recognized as an important evolutionary process that can alter gene expression patterns, including sex-biased gene expression, especially when translocations involve sex chromosomes (Wang et al. 2022). As a result, the sex chromosomes (e.g. X and Y chromosome) always enriched for sex-biased genes (Ellegren and Parsch 2007). This pattern is indeed observed in our data that female-biased genes across all tissues were significantly enriched on the X chromosome in all three seed beetles (Figure S14). To further examine the impact of gene translocation on sex-biased gene expression, we identified the derived gene traffic events between chromosomes both in *C. chinensis* (Table S7) and *C. maculatus* (Table S8) since they diverged from *A. obtectus*. In total, we identified 43 and 138 gene traffic events in *C. chinensis* and *C. maculatus*, respectively (Tables S7-S8). Of them, 35 and 107 are gene traffic between autosomes, 2 genes and 11 genes are moved from autosomes to X chromosome or vice versa, in *C. chinensis* and *C. maculatus*, respectively. We were unable to identify gene traffic events between the Y chromosome due to the rapid decay of synteny among these Bruchinae species, with very few orthologous genes can be reliably determined. This suggests that nearly all genes (97.1% in *C. chinensis* and 97.6% in *C. maculatus*) on the Y chromosome are either newly emerged or the results of translocation-based gene birth since their divergence from *A. obtectus* (Tables S9-S10).

To investigate the mechanisms underlying the translocation of genes between sex chromosomes and autosomes, we analyzed the correlation between the translocation events and the expression patterns of their corresponding orthologous genes before and after the translocation. We began by examining a case of non-replicating gene translocation, specifically the movement of a gene in *C. chinensis* from chromosome 4 to chromosome X, with the concomitant loss of the orthologous gene at the original locus (Figure 4D). Analysis of gene expression reveals that this gene is specifically highly expressed in the gonadal tissues of females, whether it is located on an autosome prior to translocation or on the X chromosome afterward, across all three species of seed beetle. Conversely, similar outcomes were observed in a gene duplication translocation event (Figure 4E). Specifically, the orthologous gene has been translocated to the Y chromosome in *C. chinensis*. Notably, prior to this translocation, the gene already demonstrates tissue-specific high expression in male gonadal tissues on the autosomes across all three seed beetles. These findings imply that the patterns of gene expression could act as a driving force in the translocation of genes. Genes that are predominantly expressed in females show a propensity to move to the X chromosome, whereas the Y chromosome serves as a preferred site for genes that are predominantly expressed in males. Furthermore, we also found that the expression profiles of certain genes shift following translocation between autosomes and sex chromosomes (Figure S15). This suggests that such translocations could potentially mitigate the effects of sexual conflict on genes.

Seed beetles exhibit numerous sexually dimorphic traits, including differences in gene expression, body size, and immunity, reflecting their evolutionary history under sexually antagonistic selection (Hotzy et al. 2012; Goenaga et al. 2015; Berg and Maklakov 2012; Berger et al. 2016; Berger et al. 2014). The ongoing recruitment of male-beneficial genes from autosomes to the Y chromosome may help partially resolve the sexual conflict between males and females to a certain extent (Kaufmann et al. 2023). We identified several genes (n=11) that exhibit changes in the direction of sex-biased expression after being translocated from autosomes to the Y chromosome in all three seed beetle species (Figure 4F). These genes were previously female-biased or unbiased in expression when located on autosomes but became male-biased after being translocated to the Y chromosome (Figure 4F). For example, *API5* (Garcia-Hughes et al. 2015), a gene essential for wing development, is located on the Y chromosome of *A. obtectus* and shows male-biased expression; whereas its ancestral copy on the autosome exhibits female-biased expression. *TDRD5*, a gene that is required for spermiogenesis in insect (Deng et al. 2022), shows strong male-biased expression when located on the Y chromosome in *A. obtectus*. We also identified genes related to metabolism (*UBQLN1* and *A1CF*) (Li et al. 2017; Blanc et al. 2021) in *C. chinensis* and a gene regulating body size in *C. maculatus* (*MTOR*) (Kaufmann et al. 2023) that exhibit male-biased expression after being translocated to the Y chromosome from autosomes. The specific expression of these gene on the Y chromosome may contribute to develop sexual dimorphism or help alleviate the sexual conflict between male and female in seed beetles.

## Discussion

The Bruchinae, as a group with significant sexual dimorphism, has its related species often used for research on sex chromosomes (Kaufmann et al. 2023; Sayadi et al. 2019; Kaufmann et al. 2021). We have successfully assembled high-quality chromosome-level genomes of three Bruchinae species: *C. chinensis*, *C. maculatus*, and *A. obtectus*, which allows us to gain deeper understanding and insight into the genomic evolution and SV within the Bruchinae family.

### Discrepancy of Y chromosome in species and populations

The assembly of Y chromosome is challenging due to the abundance of repetitive sequences on it, which complicates the process of generating a contiguous and accurate genomic sequence (Bachtrog 2013; Hughes and Page 2015). For example, the assembled Y chromosome of *C. maculatus* are relatively small compared to the Y-linked contigs reported in previous studies (Kaufmann et al. 2023). Cytological analysis across various species of seed beetles has revealed a diversity in their Y chromosome (Angus et al. 2011). Additionally, there are significant differences in Y chromosome lengths between different populations in same species, which may be attributed to widespread sexual conflict events experienced by seed beetles. This genetic variation could be a reflection of the dynamic evolutionary processes and population-specific factors influencing the structure and size of the Y chromosome in these insects. Furthermore, the substantial variation in the lengths of Y-linked contigs identified by Kaufmann *et al*. (Kaufmann et al. 2023) across different populations of the *C. maculatus* provides compelling evidence supporting the hypothesis. The sequence comparisons between the Y chromosome of the current genome of *C. maculatus* and the published Y-linked contigs suggest a higher contiguous of the current Y chromosome assembly. Furthermore, the analysis reveals that the Y-linked contig CATOUR010000029.1, as identified by Kaufmann et al. (Kaufmann et al. 2023), is actually located on chromosome 3 in our genome assembly (Figure S4E). Coverage analysis further supports that this region is part of the autosomal region, rather than the Y chromosome (Figure S4E). This observation suggests that variations in Y chromosome length across different populations could be attributed to the exchange of sequences between the Y chromosome and autosomes. Such exchanges may potentially account for the observed differences in Y chromosome length among various populations. However, these findings do not preclude the possibility that repetitive sequences may have been omitted from our assembly, potentially leading to an underestimation of the size of Y chromosome.

### Transposon amplification drives large-scale SVs in Bruchinae

Our analysis reveals distinctive SVs occurring among species within the Bruchinae family. The conservation of chromosomes in Coleoptera spans across multiple families and correlates with evolutionary distance, exhibiting a degree of decay (Van Dam et al. 2021). Similar chromosome conservation patterns were observed beyond Coleoptera, with notable instances of high-level collinearity identified across diverse families within Lepidoptera (Mathers et al. 2021). In *Drosophila*, interspecies chromosome conservation is constrained by Muller elements, which are inter-chromosomal segments that restrict recombination between different chromosomes. Comparable mechanisms have also been proposed in nematodes to elucidate chromosome conservation across diverse nematode species (Wang et al. 2022). Compared to other species within the Coleoptera order, species of the family Bruchinae have experienced extensive chromosomal recombination, resulting in complete disruption of chromosomal correspondence and showing distinctive SVs. It is noteworthy that while extensive recombination occurs among autosomes, the X chromosome exhibits relatively high conservation across the entire Coleoptera species. Similar observations have been noted in Hemiptera (Mathers et al. 2021), suggesting evolutionary constraints specific to the X chromosome (Mathers et al. 2021). Furthermore, we observed a correlation between extensive chromosomal recombination and transposable elements, particularly LINE transposons. Transposons play a crucial role as drivers of SV. Their insertion, deletion, or replication within the genome can induce functional alterations in genes or regulatory sequences, and may also prompt recombination and rearrangement of genomic regions (Cordaux and Batzer 2009; Feschotte and Pritham 2007). We observed extensive recent amplification of LINE transposons within the Bruchinae and revealed a significant correlation between these transposons and SVs between genomes. The unique amplification event of LINE transposons within the Bruchinae suggests that this may contribute to the occurrence of distinctive chromosomal variations within this taxonomic group.

### SV drive sex-biased expression of genes

Our data also indicated that SV can mitigate sexual conflict arising during evolution by modifying the gene pool of sex-biased expression. Sex-biased genes (SBGs) are not conserved among closely related species, or even among different tissues within one specie, and most SBGs are subject to certain degree of relaxed purifying selection (Tosto et al. 2023). The sex-biased expression of genes is primarily caused by intra-locus sexual conflict, which give rise to sexual dimorphism in biological traits or characteristics between males and females of a species (Tosto et al. 2023). Several studies have found a correlation between the widespread sexual conflict traits in *C. maculatus* and the large number of SBGs in its genome, and these sexual conflict traits also contribute to the unique evolutionary rate of SBGs (Kaufmann et al. 2021; Sayadi et al. 2019). Similar to *C. maculatus*, both *C. chinensis* and *A. obtectus* exhibit numerous sexual dimorphism traits between males and females (Arnqvist et al. 2022), making them excellent models for studying the evolution of SBGs and sexual conflict. Comparisons of SBGs among closely related species of Bruchinae reveal that these genes undergo rapid changes between *C. chinensis* and *C. maculatus*, which diverged 30 million years ago. The widespread differences in SBGs among closely related species of Bruchinae suggest the existence of diverse sexual dimorphism traits among them, which may have contributed to their gradual divergence from a common ancestor into distinct species (Tuda et al. 2006). Genes located at sex-conflict loci often exhibit tissue-specific and sex-biased expression patterns, making them subject to less selection pressure and thus evolving at a faster rate (Tosto et al. 2023). We found that the SBGs have a faster evolution rate and this ratio was notably elevated for those genes with stronger sex-biased expression (Figure S16), which is consistent with the prediction that SBGs should experience relaxed purifying selection (Dapper and Wade 2016). The distribution of SBGs in the genome is not completely random, and they often enriched or depleted on sex chromosomes (Ellegren and Parsch 2007). Hence, gene translocation between autosomes and sex chromosomes can effectively modify the repertoire of sex-biased expression genes, thereby influencing the degree and direction of gene expression bias and quickly adapting to sex conflicts (Ellegren and Parsch 2007).

### The effect of Y chromosome in alleviating sexual antagonism

The rapid translocation of genes on the Y chromosome provide an effective way to alleviate sexual antagonism through gene sex-specific expression. When the expression of a gene has opposite effects on different sexes, it will be subject to different selective pressures between the sexes, leading to sexual antagonism (Tosto et al. 2023). At the expression level, genes can alleviate sexual conflict at certain loci through sex-biased/specific expression, which has been proven in many species (Kaufmann et al. 2023; Pearse et al. 2019; Parrett et al. 2022). Since the Y chromosome specifically exists in male individuals, many genes that are beneficial to males but detrimental to or inconsequential for females can be transferred to the Y chromosome through autosomal translocations, thus escaping sexual selection pressure in females (Cirulis, Hansson, and Abbott 2022). Our findings suggest that genes on the Y chromosome lack synteny or orthologs across seed beetles, indicating a rapidly evolving and dynamic pattern. The collinear region between the Y chromosome and chromosome 9 of *C. chinensis* further suggests that gene translocation events between the Y chromosome and autosomes are one of the driving forces behind the emergence of new Y chromosome genes and novel sex-biased expression genes, which is also similar in *C. maculatus* (Kaufmann et al. 2023). In the seed beetles, numerous traits, including body size (Kaufmann et al. 2021), gene expression (Sayadi et al. 2019), and reproductive costs (Hotzy et al. 2012), have been discovered to vary between males and females, indicating that their genomes are subject to a great amount of sexual antagonism selection. Genes on the Y chromosome can resolve the sexual conflict between male and female traits through male-specific expression, a mechanism that has been observed on the Y chromosome of *C. maculatus* (Kaufmann et al. 2023; Sayadi et al. 2019). We identified several male-biased expression genes associated with sexual dimorphism through the Y chromosome of three types of seed beetles. However, specific functional verification requires additional experimental support.

## In conclusion

In this study, we show that three seed beetle genomes in Bruchinae undergone rapid genomic changes. The extensive genomic SVs observed in seed beetles may be associated with their rapid evolving sexual dimorphisms that causes conflicts between sexes. The high-quality reference genomes generated by this study offer an invaluable genomic resources to study the evolution of regulation of sex-biased genes and their related traits in these seed beetles.

## Method

### Study organism

The *C. chinensis* populations was collected from Shijiazhuang, Hebei province in China. The *C. maculatus* populations was collected from Beijing in China. The *A. obtectus* populations was collected from Qujing, Yunnan province in China. All insects were reared under room temperature (25-28 °C) in Beijing. *C. chinensis, C. maculatus* and *A. obtectus* were nurtured with mung bean, red bean, and white kidney bean, respectively.

### Genomic DNA isolation and sequencing

Genomic DNAs of seed beetles were extracted and purified from adult insects. DNA libraries for short-read sequencing were prepared and sequenced on the DNBseq-T7 platform (MGI Tech Co., Ltd., Shenzhen, China). Raw reads were filtered to remove low-quality bases and sequencing adaptors using Trimmomatic (v0.39) (Bolger, Lohse, and Usadel 2014). For long-read sequencing, libraries were prepared following the protocol provided for the SQK-LSK109 library preparation kit (Oxford Nanopore Technologies [ONT], Oxford, UK). DNA quantification was performed using Qubit (v4.0) (Invitrogen). Subsequently, the libraries were sequenced on an Oxford Nanopore PromethION flow cell.

### Hi-C library preparation and sequencing

Using whole bodies of adult male insects, we constructed Hi-C library of seed beetles. Briefly, genomic DNAs were extracted and purified, then digested with 100 units of *Dpn*II and marked by incubating with biotin-14-dATP. The ligated DNA was subsequently sheared into 300-600 bp fragments, followed by end-repair, A-tailing, and purification of DNA fragments. Finally, the Hi-C libraries were quantified and sequenced using the DNBseq-T7 platform (MGI Tech Co., Ltd., Shenzhen, China).

### RNA library preparation, sequencing, and data processing

Using TRIzol reagent (Thermo Fisher Scientific, Waltham, MA, USA), total mRNA of different tissues was extracted from 15 adult samples. RNA-Seq libraries were then prepared, and paired-end sequencing (at least 6G per sample) was conducted on the DNBseq-T7 platform (MGI Tech Co., Ltd., Shenzhen, China) provided by Annoroad Genomic Company. RNA sequencing from different tissues were mapped to genomes using HISAT2 (v2.2.1) (Kim et al. 2019) (--min-intronlen 20 --max-intronlen 600000 --fr -x) following SAMtools tool to generate bam files. FeatureCounts (v2.0.1) (Liao, Smyth, and Shi 2014) was used to count expression reads in each gene and reads counts between different library were normalized to transcript per kilobase per million mapped reads (TPM) for comparison. To measure expression divergence, we computed Euclidean distance following the formulas previously described (Pereira, Waxman, and Eyre-Walker 2009). RNA-Seq of *T. castaneum* are collected from public (SRR14070860, SRR14070861, SRR14070872, SRR14070873, SRR19548808, SRR19548809, SRR19548810, SRR19548811).

### Genome assembly and annotation

De novo assembly with NextDenovo (v2.4.0) (Jiang et al. 2023) using default setting gave the initial assembly of contigs. The NextDenovo assembly was polished with two round of Pilon (v1.24) (Walker et al. 2014) using 150-bp paired-end reads (∼162, 134 and 292 coverage of the *C. chinensis*, *C. maculatus*, and *A. obtectus* genomes). For the highly heterozygous seed beetles’ genomes, we use purge_haplotigs(Roach, Schmidt, and Borneman 2018) to reduce its heterozygosity. Finally, we used Hi-C data (∼224, 293 and 76 coverage of the *C. chinensis*, *C. maculatus*, and *A. obtectus* genomes) and assembled the polished contigs into chromosomes using the standard process of the ALLHIC (v0.9.8) (Zhang et al. 2019). We visualized the hic matrix with Juicebox (v1.11.08) (Durand et al. 2016) and adjusted the order and orientation of contigs manually.

RepeatModeler (v2.0.1) (Flynn et al. 2020) with the default parameters was used to generated the custom TE libraries of three genomes. The repetitive sequence in genomes were annotated by RepeatMasker (v4.0.9) (http://www.repeatmasker.org) using custom TE libraries. To gain deeper insights into the evolutionary history of the TE sequences, we estimated their insertion times in three seed beetles using a Kimura distance-based analysis with the parseRM pipeline.

For de novo prediction, we utilized RNA-Seq alignments in BAM files to train the AUGUSTUS (v3.4.0) (Stanke et al. 2006) gene prediction tool via BRAKER (v2.1.4) (Hoff et al. 2016). Importantly, we imported the protein-coding sequences of *C. chinensis*, *C. maculatus*, and *A. obtectus* into miniport (v0.11) (Li 2023) with the parameters “-I600kb --gff” to generate gene structures based on orthologous evidence. In the transcriptome-based prediction, raw reads underwent filtering by Trimmomatic (v0.39). These filtered reads were subsequently aligned to the genome assembly using HISAT2 (v2.2.1) (Kim et al. 2019) to determine transcript positions and extract transcript sequences using StringTie (v2.2.1) (Pertea et al. 2015). Next, we mapped the extracted transcript sequences to the genomes using PASA (v2.4.1) (Haas et al. 2008). Additionally, TransDecoder (v5.5.0) (https://github.com/TransDecoder/TransDecoder) was employed to generate gene predictions using PASA-extracted transcripts. Finally, we integrated gene predictions from the three methods using EVidenceModeler (v2.1.0), assigning weights for each method (Augustus: 2; TransDecoder: 3; Miniport: 8; PASA: 10;). The parameters used were: “--segmentSize 1000000 --overlapSize 100000”.

### Identification of orthologous proteins

To maximize the identification of orthologous protein pairs between species, we employed BLAST to search for sequence relationships between proteins from two species. For each gene alignment, we selected the orthologous gene with the lowest E-value in the comparison, using one species as a reference.

To identify orthologous protein relationships between the Y chromosome and autosomes within the species, we utilized the Y chromosome protein sequence as a reference. Using BLAST, we searched for autosomal protein with the lowest E-value to the Y-linked protein and subsequently calculated the sequence identity between them.

### Phylogenetic tree construction and species divergence time estimation

Eight coleopteran species (*Pachyrhynchus sulphureomaculatus*, *Pyrochroa serraticornis*, *T. castaneum*, *Propylea japonica*, *Rhagonycha fulva*, *Pogonus chalceus*, *Tribolium confusum* and *Tribolium freemani*) and our three genomes with *D. melanogaster* as outgroup were selected, and their longest transcripts were extracted for orthologs inference using OrthoFinder (v2.5.4) (Emms and Kelly 2019). Subsequently, MAFFT (v7.508) (Katoh and Standley 2013) was employed for sequence alignment of the resulting single-copy orthologous genes. Gblocks (v0.91b) (Castresana 2000) was utilized to extract conserved sites with parameters ‘-b4=5 -b5=h -t=p’ for phylogenetic tree inference. The best-fitting model (Q. insect+F+I+I+R5) was determined by the ModelFinder program implemented in IQ-TREE (v2.2.0.3) (Nguyen et al. 2015) based on the Bayesian information criterion and ultrafast bootstrap (Hoang et al. 2018) approximation with 1000 replicates. Divergence time was estimated using MCMCTree, a tool within the PAML (v4.9) (Yang 2007) package. Calibration information of fossil nodes was obtained from the Timetree website.

### Coverage and synteny analyses

We mapped whole genome sequencing reads from female and male individuals to the assemblies using the aln function of bwa (v0.7.17-r1188). Alignments were filtered for uniquely mapped reads and average chromosomes coverage was calculated using bedtools with 100-kb windows. M:F-fold change in coverage of each chromosomes was calculated to determine the X/Y chromosome.

To examine the degree of collinearity between species, we used MCscanX (Wang et al. 2012) to identify collinearity regions between different genomes. The syntenic blocks are defined as regions with a similar order of gene distribution in different genomes, comprising at least 5 genes (-s 5). We extracted 1367 BUSCO loci (insect v10) coordinates to calculated the gene order conservation (GOC) score using a common script (Van Dam et al. 2021). The orthologous proteins identity (OPI) was calculated by average BUSCO loci similarity between genomes.

### Identification of evolutionary chromosomal structural variants

Pairwise genome alignments were performed between different genomes, respectively, using mummer (v4.0.0rcl) (Marçais et al. 2018). Sequencing similarity of collinearity regions were calculated by nucmer with custom parameters (--mum -l 10 -c 100 -d 10). The resulting alignments were then as the input for SyRI (v1.6) (Goel et al. 2019) to identify SVs. The SVs with a size < 5k were filtered manually and the boundary of SVs were considered as breakpoints. The subroutine plotsr (-H 8 -W 5) of SyRI is used for visualization.

### Identification and evolutionary analysis of sex-biased genes

DESeq2 (v1.38.3) (Love, Huber, and Anders 2014) was used to estimate differential gene expression between the sexes in different tissue samples. Gene with low read counts (= 0) in any samples would be filtered and the read counts for the remain gene were normalized on a log2 scale. Genes exhibiting fold change ≥ 1.5 in male-to-female expression were categorized as male-biased genes, while genes with fold change ≥ 1.5 in female-to-male expression were categorized as female-biased genes.

To visualize the impact of gene bias pattern on gene evolution rate, we assigned each pair of single-copy orthologous genes to one of seven bins of predefined ranges of log2FC in gene expression: extreme female bias (log2FC < −1), strong female bias (−1 ≤ log2FC < −0.59), moderate female bias (−0.59 ≤ log2FC < 0), moderate male bias (0 ≤ log2FC < 0.59), strong male bias (0.59 ≤ log2FC < 1), and extreme male bias (log2FC ≥ 1). Only gene families containing orthologous genes from both *A. obtectus* and *C. chinensis* / *C. maculatus* within the same box were assessed for their evolutionary rates from *A. obtectus* to *C. chinensis* / *C. maculatus*. Categorization was carried out separately based on expression levels in heads, gut and gonads. Evolutionary rate of orthologous sex-biased genes was calculated respectively using ParaAT (v1.0) (Zhang et al. 2012) with default parameters.

### Transcriptomic heterozygosity analysis

Variant was called from bam files of mapped RNA-Seq reads by a subroutine HaplotypeCaller of the Genome Analysis Toolkit UnifiedGenotyper (GATK) (v3.8) (McKenna et al. 2010). Variants were subsequently filtered to exclude those with Fisher Strand values exceeding 20.0 or Qual-By-Depth values below 3.0 using VCFtools (v0.1.16) (Danecek et al. 2011). Finally, we used a custom script to summarize the number of heterozygous and homozygous SNP loci on each chromosome.

### X Chromosomal Enrichment Analysis

To identity the enrichment of sex-biased genes on the X chromosome of seed beetles, we used a method previously described (Catalán, Macias-Muñoz, and Briscoe 2018). Briefly, we counted the sex-biased genes on the X chromosome and compared them with the autosomes. Chromosomal counts of sex-biased and unbiased genes were tabulated, and a 2×2 contingency table was constructed to facilitate Fisher’s exact test. This test was employed to assess deviations in the proportions of sex-biased and unbiased genes between the X-chromosome and autosomes.

### Gene translocation analysis

To identify the gene translocation events, we first use BLAST with the lowest E-value to search for orthologous genes between species. Next, we employ MCscanX to identify collinear genes across species. It is hypothesized that species-specific gene translocation events will not exhibit collinearity with other species, whereas the orthologous genes among the remaining species should maintain collinearity. Additionally, we utilized custom script to exclude gene translocation events that were incorrectly identified due to tandem repeats. Finally, we manually filtered the remaining genes and obtained accurate gene translocation events.

## Acknowledgements

This work was supported by National Key R*&*D Program of China (2021YFC2600100), the Open Research Fund Program of State Key Laboratory of Integrated Management of Pest Insects and Rodents (Grant No. IPM2201), the National Natural Science Foundation of China (Nos. 31571737), and Initiative Scientific Research Program, Institute of Zoology, Chinese Academy of Sciences (2023IOZ0203).

## Author contributions

X.Z., Y.L., and J.C. conceived and designed the study. M.L., C.L., and X.W. reared the seed beetles and performed experiments. W.C., J.Z., and L.X. assembled the genome, annotated protein-coding genes and TEs, and performed data analyses. W.C., J.Z., X.Z., Y.L., and J.C. wrote and revised the manuscript.

## Data availability

The raw sequences of Nanopore ultra-long reads, WGS short reads, RNA-seq reads, and Hi-C reads have been deposited in the National Center for Biotechnology Information SRA (BioProject accession no. PRJNA792679 for *C. chinensis* and *C. maculatus* and PRJNA912403 for *A. obtectus*). The genome assembly has been deposited in the National Center for Biotechnology Information Genome (JBICBS000000000 for *C. chinensis*; JBCIKN000000000 for *C. maculatus* and JBHEVA000000000 for *A. obtectus*). The assembled genome sequences and gene and transposable element annotations and the scripts used for analyses are available on Zenodo (https://doi.org/10.5281/zenodo.13744296). All study data are included in the main article and Supplementary Materials.

## Code availability

All codes or tools used in this study are described in the methods and available on Zenodo (https://doi.org/10.5281/zenodo.13744296).

## Supplementary materials

**Figure S1. Genomic characteristics of three seed beetles.**

**Figure S2. Comparison of *C. maculatus* genome assembly with previously published *C. maculatus* genomes.**

**Figure S3. Independent Hi-C matrix and sequence alignment of X chromosomes.**

**Figure S4. Independent Hi-C matrix and sequence alignment of Y chromosomes.**

**Figure S5. Analysis of collinearity between Coleoptera species.**

**Figure S6. Sequence differences between the genomes of seed beetles.**

**Figure S7. Insertion history of TEs among Coleoptera species.**

**Figure S8. Chromosomal conservation of orthologous genes between Bruchinae and *P. sulphureomaculatus*.**

**Figure S9. Allelic heterozygosity in *C. chinensis’* transcriptome.**

**Figure S10. Dose expression pattern of sex chromosomes for seed beetles.**

**Figure S11. The evolutionary rate of Y-linked genes in three seed beetles.**

**Figure S12. Comparison of sex-biased pattern of orthologous genes in three seed beetles.**

**Figure S13. Boxplot for the density of classification genes inside/outside SV.**

**Figure S14. X Chromosomal enrichment analysis of sex-biased genes.**

**Figure S15. Examples of gene transfer resulting in genetic environmental changes and altered sex bias in gene expression.**

**Figure S16. The evolutionary rate of sex-biased genes.**

**Table S1. Assembly statistics of three seed beetle genomes**

**Table S2. Transposable element annotation of *Callosobruchus chinensis***

**Table S3. Transposable element annotation of *Callosobruchus maculatus***

**Table S4. Transposable element annotation of *Acanthoscelides obtectus***

**Table S5. Identity of the single-copy protein similarity matrix for 12 species (n=1397)**

**Table S6. Gene Order Conservations score matrix for 12 species (n=1397)**

**Table S7. Summary of gene traffic between *A. obtectus* and *C. chinensis***

**Table S8. Summary of gene traffic between *A. obtectus* and *C. maculatus***

**Table S9. Summary of *C. chinensis* Y-links gene traffic compared to *A. obtectus***

**Table S10. Summary of *C. maculatus* Y-links gene traffic compared to *A. obtectus***

